# Network Enhancement: a general method to denoise weighted biological networks

**DOI:** 10.1101/317941

**Authors:** Bo Wang, Armin Pourshafeie, Marinka Zitnik, Junjie Zhu, Carlos D. Bustamante, Serafim Batzoglou, Jure Leskovec

**Author notes:** These authors contributed equally (Bo Wang, Armin Pourshafeie, Marinka Zitnik).

## Abstract

Networks are ubiquitous in biology where they encode connectivity patterns at all scales of organization, from molecular to the biome. However, biological networks are noisy due to the limitations of technology used to generate them as well as inherent variation within samples. The presence of high levels of noise can hamper discovery of patterns and dynamics encapsulated by these networks. Here we propose Network Enhancement (NE), a novel method for improving the signal-to-noise ratio of undirected, weighted networks, and thereby improving the performance of downstream analysis. NE applies a novel operator that induces sparsity and leverages higher-order network structures to remove weak edges and enhance real connections. This iterative approach has a closed-form solution at convergence with desirable performance properties. We demonstrate the effectiveness of NE in denoising biological networks for several challenging yet important problems. Our experiments show that NE improves gene function prediction by denoising interaction networks from 22 human tissues. Further, we use NE to interpret noisy Hi-C contact maps from the human genome and demonstrate its utility across varying degrees of data quality. Finally, when applied to fine-grained species identification, NE outperforms alternative approaches by a significant margin. Taken together, our results indicate that NE is widely applicable for denoising weighted biological networks, especially when they contain high levels of noise.

---

Networks provide an elegant abstraction for expressing fine-grained connectivity and dynamics of interactions in complex biological systems ^1^. In this representation, the nodes indicate the components of the system. These nodes are often connected by non-negative, (weighted-)edges which indicate the similarity between two components. For example, in protein-protein interaction (PPI) networks, weighted edges capture the strength of physical interactions between proteins and can be leveraged to detect functional modules ^2^. However, accurate experimental quantification of interaction strength is challenging ^3, 4^. Technical and biological noise can lead to superficially strong edges, implying spurious interactions; conversely, dubiously weak edges can hide real, biologically important connections ^4–6^. Furthermore, corruption of experimentally derived networks by noise can alter the entire structure of the network by modifying the strength of edges within and amongst underlying biological pathways. These modifications adversely impact the performance of downstream analysis ^7^. The challenge of noisy interaction measurements is not unique to PPI networks and plagues many different types of biological networks, such as Hi-C ^8^ and cell-cell interaction networks ^9^.

To overcome this challenge, computational approaches have been proposed for denoising networks. These methods operate by replacing the original edge weights with weights obtained based on a diffusion defined on the network ^10, 11^. However, these methods are often not tested on different types of networks ^11^, rely on heuristics without providing explanations for why these approaches work, and lack mathematical understanding of the properties of the denoised networks ^10, 11^. Consequently, these methods may not be effective on new applications derived from emerging experimental biotechnology.

Here, we introduce Network Enhancement (NE), a novel diffusion-based algorithm for network denoising that does not require supervision or any prior knowledge. NE takes as input a noisy, undirected, weighted network and outputs a network on the same set of nodes but with a new set of edge weights (Figure 1). The main crux of NE is the observation that nodes connected through paths of high-weight edges are more likely to have a direct, high-weight edge between them ^12, 13^. Following this intuition, we define a diffusion process that uses random walks of length three or less and a form of regularized information flow to denoise the input network (Figure 1A and **Methods**). Intuitively, this diffusion generates a network in which nodes with strong similarity/interactions are connected by high-weight edges while nodes with weak similarity/interactions are connected by low-weight edges (Figure 1B). Mathematically, this means that eigenvectors associated with the input network are preserved while the eigengap is increased. In particular, NE denoises the input by down-weighting small eigenvalues more aggressively than large eigenvalues. This re-weighting is advantageous when the noise is spread in the eigen-directions corresponding to small eigenvalues ^14^. Furthermore, the increased eigengap of the enhanced network is a highly appealing property as it leads to accurate detection of modules/clusters ^15, 16^ and allows for higher-order network analysis ^12^. Moreover, NE has an efficient and easy to implement closed-form solution for the diffusion process and provides mathematical guarantees for this converged solution. (Figure 1B and **Methods**).

**Figure 1:**
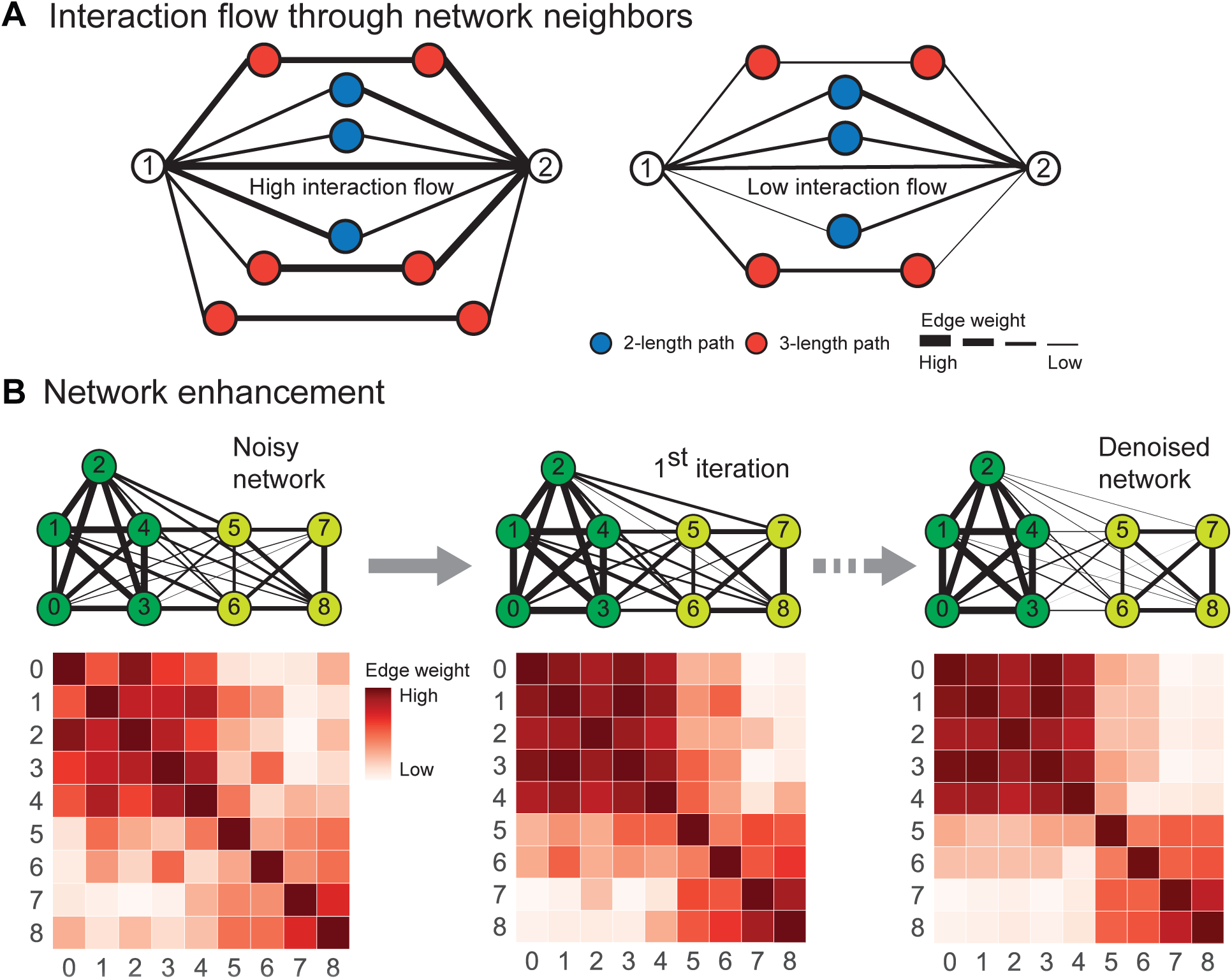
Overview of Network Enhancement (NE). (**A**) NE employes higher-order network structures to enhance a given weighted biological network. The diffusion process in NE revises edge weights in the network based on interaction flow between any two nodes. Specifically, for any two nodes, NE updates the weight of their edge by modeling all paths of length three or less connecting those nodes. (**B**) The iterative process of NE. NE takes as input a weighted network and the associated adjacency matrix (visualized as a heat map). It then iteratively updates the network using the NE diffusion process, which is guaranteed to converge. The diffusion defined by NE improves the input network by strengthening edges that are close to other strong edges in the network according to NE’s diffusion distance or are supported by many weak edges. On the other hand, NE weakens edges that are not supported by many strong edges. Upon convergence, the enhanced network is a symmetric, doubly stochastic matrix (DSM) (**Supplementary Note 2**), which makes the enhanced network well-suited for downstream computational analysis. Furthermore, through enforcement of the DSM structure NE produces sparse networks with lower noise levels.

## Results

We have applied NE to three challenging yet important problems in network biology. In each experiment, we evaluate the network denoised by NE against the same network denoised by alternative methods: network deconvolution (ND) ^10^ and diffusion state distance (DSD) ^11^. For completeness, we also compare our results to a network reconstructed from features learned by Mashup (MU) ^17^. All three of these methods use a diffusion process as a fundamental step in their algorithms and have a closed-form solution at convergence. ND solves an inverse diffusion process to remove the transitive edges, and DSD uses a diffusion-based distance to transform the network. While ND and DSD are denoising algorithms, Mashup is a feature learning algorithm that learns low-dimensional representations for nodes based on their steady-state topological positions in the network. This representation can be used as input to any subsequent prediction model. In particular, a denoised network can be constructed by computing a similarity measure using MU’s output features ^17^.

### NE improves human tissue networks for gene function prediction

Networks play a critical role in capturing molecular aspects of precision medicine, particularly those related to gene function and functional implications of gene mutation ^18, 19^. We test the utility of our denoising algorithm in improving gene interaction networks from 22 human tissues assembled by Greene *et al.* ^20^. These networks capture gene interactions that are specific to human tissues and cell lineages ranging from B lymphocyte to skeletal muscle and the whole brain ^20, 21^. We predict cellular functions of genes specialized in different tissues based on the networks obtained from different denoising algorithms.

Given a tissue and the associated tissue-specific gene interaction network, we first denoise the network and then use a network-based algorithm on the denoised edge weights to predict gene functions in that tissue. We use standard weighted random walks with restarts to propagate gene-function associations from training nodes to the rest of the network ^22^. We define a weighted random walk starting from nodes representing known genes associated with a given function. At each time step, the walk moves from the current node to a neighboring node selected with a probability that depends on the edge weights and has a small probability of returning to the initial nodes ^22^. The algorithm scores each gene according to its visitation probability by the random walk. Node scores returned by the algorithm are then used to predict gene-function associations for genes in the test set. Predictions are evaluated against experimentally validated gene-function associations using a leave-one-out cross-validation strategy.

When averaged over the four denoising algorithms and the 22 human tissues, the gene function prediction improved by 12.0% after denoising. Furthermore, we observed that all denoising algorithms improved the average prediction performance (Figure 2A and **Supplementary Data**). These findings motivate the use of denoised networks over original (raw) biological networks for downstream predictive analytics. We further observed that gene function prediction performed consistently better in combination with networks revised by NE than in combination with networks revised by other algorithms. On average, NE outperformed networks reconstructed by ND, DSD, and MU by 12.3%. In particular, NE resulted in an average a 5.1% performance gain over the second best-performing denoised network (constructed by MU). Following Greene *et al.* ^20^, we further validated our network enhancement approach by examining each enhanced tissue network in turn and evaluating how well relevant tissue-specific gene functions are connected in the network. The expectation is that function-associated genes tend to interact more frequently in tissues in which the function is active than in other non-relevant tissues ^20^. As a result, relevant functions are expected to be more tightly connected in the tissue network than functions specific to other tissues. For each NE-enhanced tissue network, we ranked all functions by the edge density of function-associated tissue subnetworks and examined top-ranked functions. In the NE-enhanced blood plasma network, we found that functions with the highest edge density were blood coagulation, fibrin clot formation, and negative regulation of very-low-density lipoprotein particle remodeling, all these functions are specific to blood plasma tissue (Figure 2B). This finding suggests that tissue subnetworks associated with relevant functions tend to be more connected in the tissue network than subnetworks of non-tissue-specific functions. The most connected functions in the NE-enhanced brain network were brain morphogenesis and forebrain regionalization, which are both specific to brain tissue (Figure 2B). Examining edge density-based rankings of gene functions across 22 tissue networks, we found relevant functions consistently placed at or near the top of the rankings, further indicating that NE can improve signal-to-noise ratio of tissue networks.

**Figure 2:**
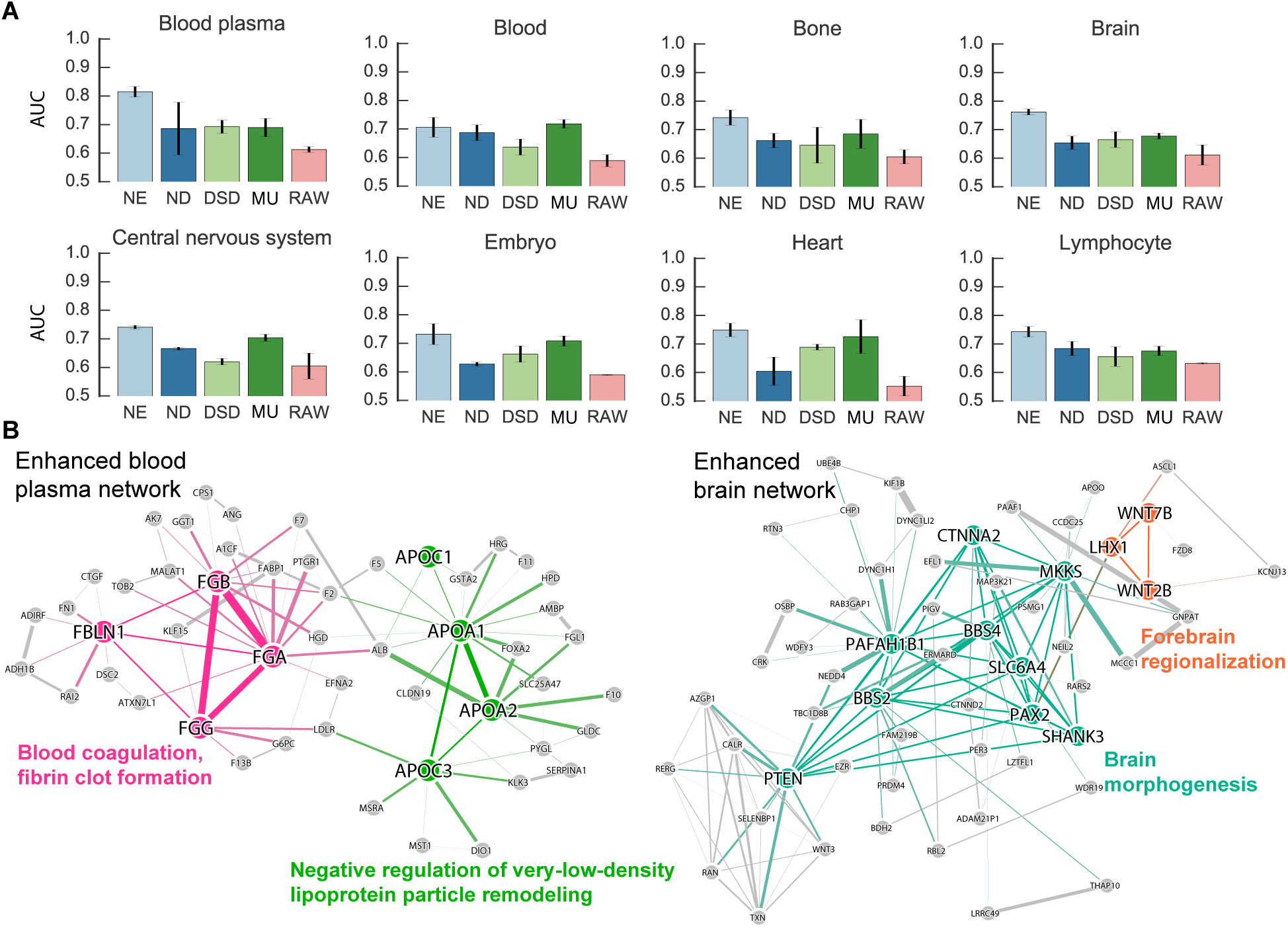
Gene function prediction for genome-wide gene interaction networks in human tissues. (**A**) We assessed the utility of original networks (RAW) and networks denoised using MU, ND, DSD and NE for tissue-specific gene function prediction. Each bar indicates the performance of a network-based approach that was applied to a raw or denoised gene interaction network in a particular tissue and then used to predict gene functions in that tissue. Prediction performance is measured using the area under receiver operating characteristic curve (AUROC), where a high AUROC value indicates the approach learned from the network to rank an actual association between a gene and a tissue-specific function higher than a random gene, tissue-specific function pair. Error bars indicate performance variation across tissue-specific gene functions. Results are shown for eight human tissues, additional fourteen tissues are considered in **Supplementary Figures 1 and 2**. (**B**) For blood plasma and brain tissues, we show gene interaction subnetworks centered on two blood plasma gene functions and two brain gene functions with the highest edge density in NE-denoised data. Edge density for each gene function (with *n* associated genes) was calculated as the sum of edge weights in the NE-denoised network divided by the total number of possible edges between genes associated with that function (*n* × (*n* − 1)/2). The most interconnected gene functions in each tissue (shown in color, names of associated genes are emphasized), are also relevant to that tissue.

### NE improves Hi-C interaction networks for domain identification

The recent discovery of numerous cis-regulatory elements away from their target genes points to the deep impact of 3D structure of DNA on cell regulation and reproduction ^23–25^. Chromosome conformation capture (3C) based technologies ^25^ provide experimental approaches for understanding the chromatin interactions within DNA. Hi-C is a 3C-based technology that allows measurement of pairwise chromatin interaction frequencies within a cell population ^8, 25^. The Hi-C reads are grouped into bins based on the genomic region, where the bin size determines the measurement resolution.

Hi-C read data can be thought of as a network where genomic regions are nodes and the normalized read counts mapped to two bins are weighted edges. To identify clusters of genomic regions that are close in the 3D genome structure we can use network community detection algorithms with the Hi-C derived networks ^26^. The detected megabase-scale communities correspond to regions known as topological associating domains (TADs) and represent chromatin interaction neighborhoods ^26^. TADs tend to be enriched for regulatory features ^27, 28^ and are hypothesized to specify elementary regulatory micro-environment. Therefore, detection of these domains is important for the analysis and interpretation of Hi-C data. The limited number of Hi-C reads, the hierarchical structure of TADs and other technological challenges lead to noisy Hi-C networks, and can hamper accurate detection of TADs ^25^.

To investigate the ability of NE to improve TAD detection, we apply NE to a Hi-C dataset and analyze the performance of a standard domain identification pipeline with and without a network denoising step. We used 1kb and 5kb resolution Hi-C data from all autosomes of the GM12878 cell line ^8^. Since true gold-standards for TAD regions are lacking, the true cluster assignments were determined as detailed in **Supplementary Data**. Figure 3A shows a heatmap of the raw Hi-C data for a portion of chromosome 16.

**Figure 3:**
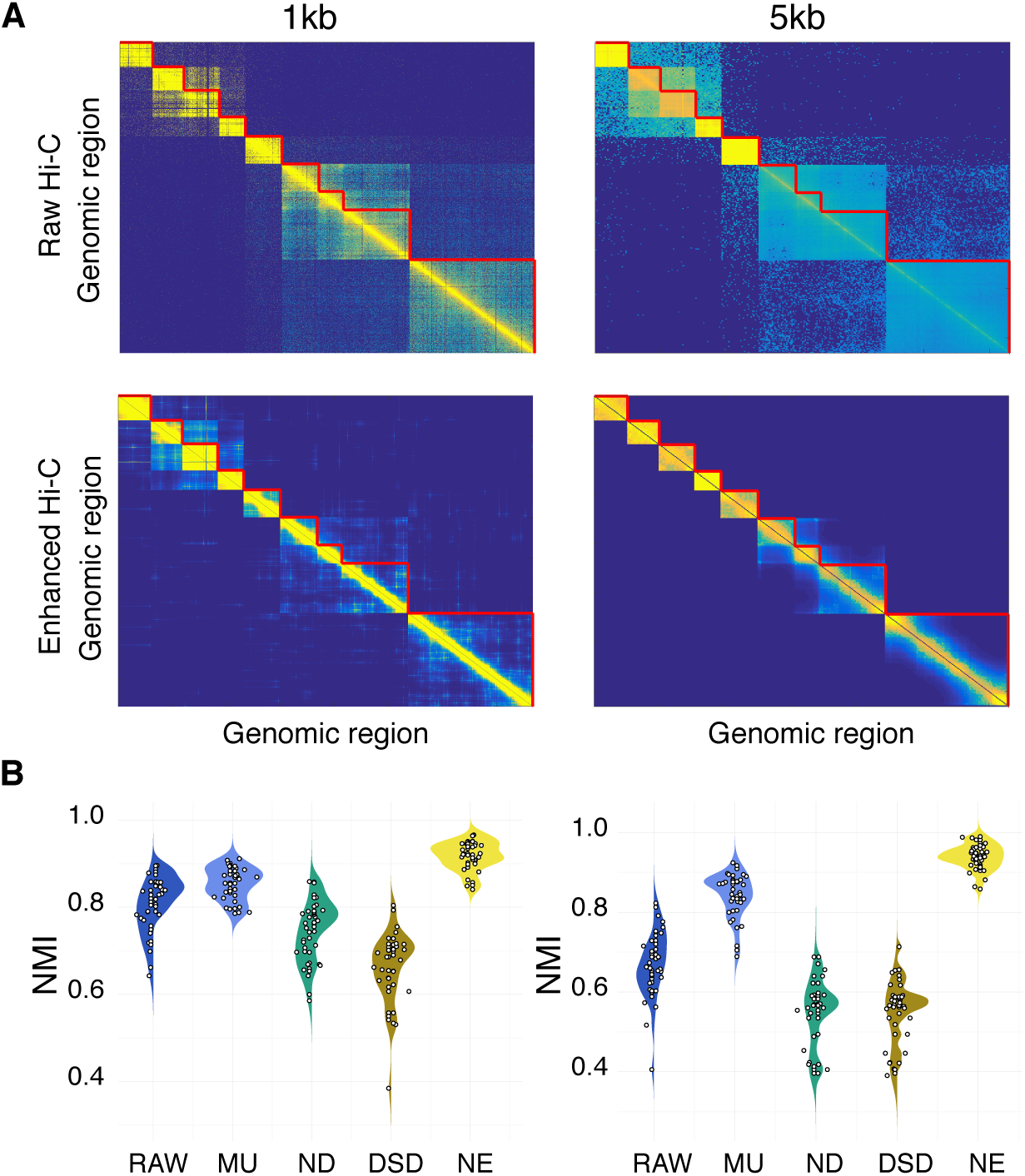
Domain identification in Hi-C genomic interaction networks. (**A**) Heatmap of Hi-C contact matrix for a portion of chromosome 16. When 1kb resolution data is denoised with NE and Louvain community detection (**S**upplementary Data) is used, the NMI of chromosome 16 corresponds to the median, therefore, this chromosome was chosen as a fair representation of the overall performance. The top two heatmaps show the contact matrices for original (raw) data and the bottom heatmaps represent the contact matrices for data after application of NE. The images on the left correspond to data with 1kb resolution (*i.e.*, each bin contains a 1kb region) and the right images correspond to the same section at 5kb resolution. The red lines indicate the borders for each domain as detailed in **Supplementary Data**. In each case, the network is consisted of genomic windows of length 1kb (left) or 5kb (right) as nodes, and normalized number of reads mapped to both regions as edge weights. The data was truncated for visualization purposes. (**B**) NMI for clusters detected. For each algorithm, the left side of the violin plot corresponds to Louvain community detection algorithm and the right side corresponds to MSCD algorithm. Each dot indicates the performance on a single autosome (The distance of the dots from the vertical axis is dictated by noise and is for visualization only). While for raw data and data preprocessed with DSD and ND the overall NMI decreases as resolution decreases, for NE and MU the performance remains high. MU maintains good overall performance with lower resolution, however, the increase in the spread of the data indicates that the consistency of performance has decreased compared to NE where the spread of NMI remains the same.

We applied two community detection methods (**Supplementary Data**) to each Hi-C network and compared the quality of the detected TADs with or without network denoising. Visual inspection of the Hi-C contact matrix before and after Hi-C network is denoised using NE reveals an enhancement of edges within each community and sharper boundaries between communities (Figure 3A). This improvement is particularly clear for the 5kb resolution data, where communities that were visually undetectable in the raw data become clear after denoising with NE. To quantify this enhancement, the communities obtained from raw networks, as well as, networks enhanced by NE or other denoising methods were compared to the true cluster assignments. We used normalized mutual information (NMI, **Supplementary Note 1**) as a measure of shared information between the detected communities and the true clusters. NMI ranges between 0 to 1, where a higher value indicates higher concordance and 1 indicates an exact match between the detected communities and the true clusters. The results across 22 autosomes indicate that while denoising can improve the detection of communities, not all denoising algorithms succeed in this task (Figure 3B). For both resolutions considered, NE performs the best with an average NMI of 0.92 for 1kb resolution and 0.94 for 5kb resolution, MU (the second best performing method) achieves an average NMI of 0.85 and 0.84, respectively while ND and DSD achieve lower average NMI than the raw data which has NMI of 0.81 and 0.67, respectively. Furthermore, we note that the performance of NE and MU remains high as the resolution decreases from 1kb to 5kb, in contrast, the ability of the other pipelines in detecting the correct communities diminishes. While MU maintains a good performance at 5kb resolution, due to relatively poor performances on a few chromosomes, the standard deviation of NMI values after denoising with MU increases from 0.037 in 1kb data to 0.054 in 5kb data. On the other hand, the NMI values for data denoised with NE maintain a similar spread for both resolutions (standard deviation 0.033 and 0.031, respectively). The better average NMI and smaller spread indicate that NE can reliably enhance the network and improve TAD detection.

### NE improves similarity network for fine-grained species identification

Fine-grained species identification from images concerns querying objects within the same subordinate category. Traditional image retrieval works on high-level categories (*e.g.*, finding all butterflies instead of cats in a database given a query of a butterfly), while fine-grained image retrieval aims to distinguish categories with subtle differences (*e.g.*, monarch butterfly versus peacock butterfly). One major obstacle in fine-grained species identification is the high similarity between subordinate categories. On one hand, two subordinate categories share similar shapes and carry the subtle color difference in a small region; on the other hand, two subordinate categories of close colors can only be well separated by texture. Furthermore, viewpoint, scale variation, and occlusions among objects all contribute to the difficulties in this task ^29^. Due to these challenges, similarity networks, which represent pair-wise affinity between images, can be very noisy and ineffective in the retrieval of a sample from the correct species for any query.

We test our method on the Leeds butterfly fine-grained species image dataset ^30^. Leeds Butterfly dataset contains 832 butterflies in 10 different classes with each class containing between 55 to 100 images ^30^. We use two different common encoding methods (Fisher Vector (FV) and Vector of Linearly Aggregated Descriptors (VLAD) with dense SIFT; **Supplementary Data**) to generate two different vectorizations of each image. These two encoding methods describe the content of the images differently and therefore can contain different information. Each descriptor can generate a similarity network in which nodes represent images while edge weights indicate similarity between pairs of images. The inner product of these two similarity networks is used as the single input network to other denoising algorithms.

Visual inspection indicates that NE is able to greatly improve the overall similarity network for fine-grain identification (Figure 4A). While both encodings partially separate the species, before applying NE, all the images are tangled together without a clear clustering. On the other hand, the resulting similarity network after applying NE clearly shows 10 clusters corresponding to different butterfly species (Figure 4A). More specifically, given a certain query, the original input networks fail to capture the true affinities between the query butterfly and its most similar retrievals, while NE is able to correct the affinities and more reliably output the correct retrievals (Figure 4B).

**Figure 4:**
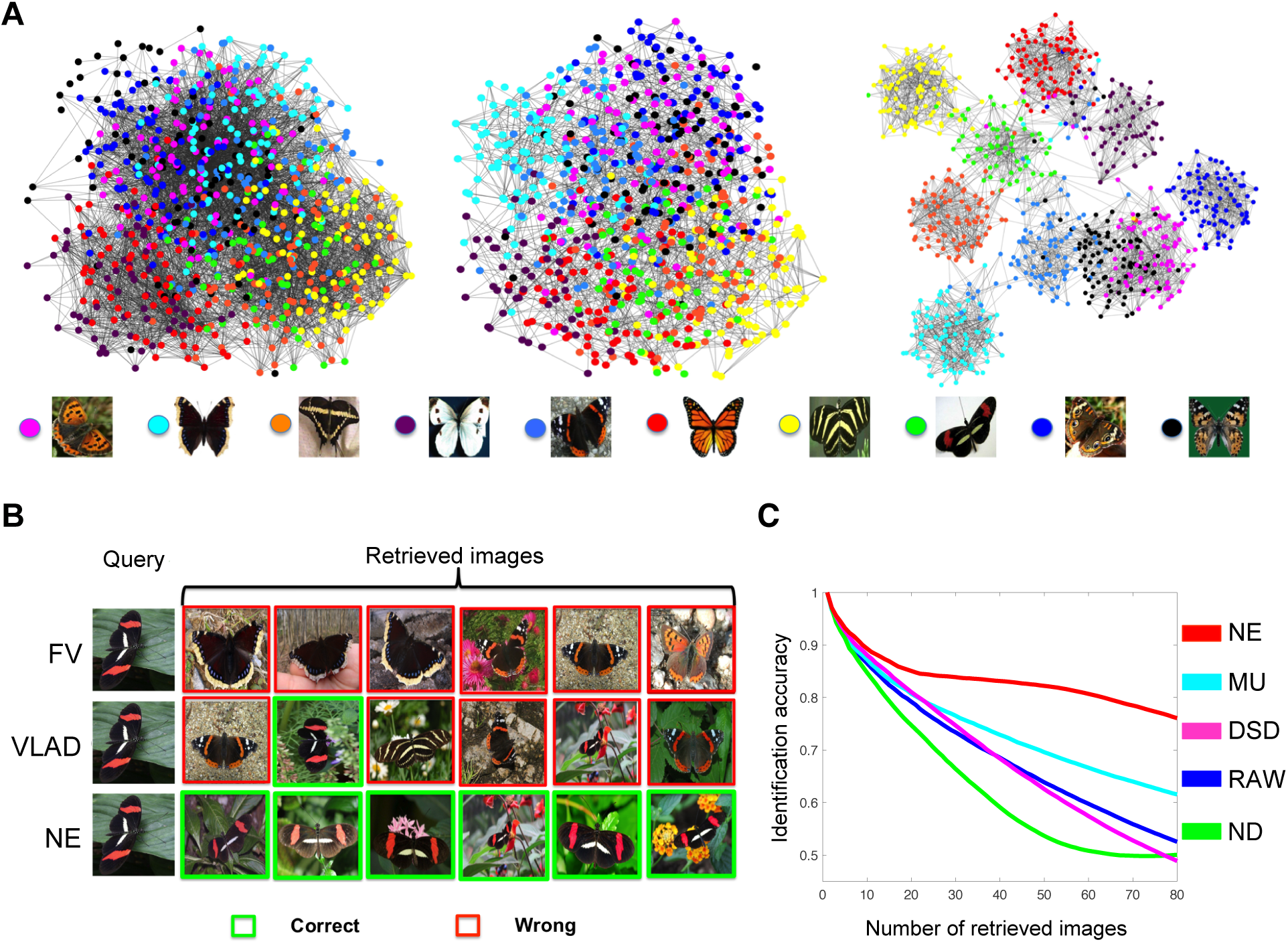
Network-based butterfly species identification. (Best seen in color.) Example of combining different metrics to improve the retrieval performance. (**A**) Visualization of encoded images as a network. From left to right: Fisher Vector, VLAD (**Supplementary Data**), and the denoised similarity network by our method (NE). The legend shows an example of each species included in the network. (**B**) Retrieval by each encoding method. Given a query butterfly, original descriptors fail to retrieve the correct species while the network denoised by NE is able to recover the correct similarities between the query and its neighbors within the same class. (**C**) Species identification accuracy when varying the number of retrieved images. A detailed comparison with other methods. The curves show the identification accuracy (**Supplementary Note 1**) as a function of number of retrievals.

To quantify the improvements due to NE in the task of species identification, we use identification accuracy, a standard metric which quantifies the average numbers of correct retrievals given any query of interests (**Supplementary Note 1**). A detailed comparison between NE and other alternatives by examining identification accuracy of the final network with respect to a different number of top retrievals demonstrates NE’s ability in improving the original noisy networks (Figure 4B). For example, when considering top 40 retrievals, NE can improve the raw network by 18.6% (more than 10% better than other alternatives). Further, NE generates the most significant improvement in performance (41% over the raw network and more than 25% over the second best alternative), when examining the top 80 retrieved images.

Current denoising methods suffer from high sensitivity to the hyper-parameters when constructing the input similarity networks, e.g., the variance used in Gaussian kernel (**Supplementary Note 1**). However, our model is more robust to the choice of hyper-parameters (**Supplementary Figure 3**). This robustness is due to the strict structure enforced by the preservation of symmetry and DSM structure during the diffusion process (see **Supplementary Note 2**).

## Discussion

We proposed Network Enhancement as a general method to denoise weighted undirected networks. NE implements a dynamic diffusion process that uses both local and global network structures to construct a denoised network from its noisy version. The core of our approach is symmetric, positive semi-definite, doubly stochastic matrix, which is a theoretically justified replacement for the commonly used row-normalized transition matrix ^31^. We showed that NE’s diffusion model preserves the eigenvectors and increases the eigengap of this matrix for large eigenvalues. This finding provides insight into the mechanism of NE’s diffusion and explains its ability to improve network quality. ^15, 16^ In addition to increasing the eigengap, NE disproportionately trims small eigenvalues. This property can be contrasted with the principal component analysis (PCA) where the eigenspectrum is truncated at a particular threshold. Through extensive experimentation, we show that NE can flexibly fit into important network analytic pipelines in biology and that its theoretical properties enable substantial improvements in the performance of downstream network analyses.

We see many opportunities to improve upon the foundational concept of NE in future work. First, in some cases, a small subset of high confidence nodes may be available. For example, genomic regions in the Hi-C contact maps can be augmented using data obtained from 3C technology or a small number of species can be identified by a domain expert and used together with network data as input to a denoising methodology. Extending NE to take advantage of the small amount of accurately labeled data might further extend our ability to denoise networks. Second, although we showed the utility of NE for denoising several types of weighted networks, there are other network types worth exploring, such as multimodal networks involving multi-omic measurements of cancer patients. Finally, incorporating NE’s diffusion process into other network analytic pipelines can potentially improve the performance of these pipelines. For example, Mashup ^17^ learns vector representations for nodes based on a steady state of a traditional random walk with restart, and replacing Mashup’s diffusion process with the rescaled steady state of NE might be a promising future direction.

## Methods

### Problem definition and doubly stochastic matrix property

Let *G* = (*E*, *V*, *W*) be a weighted network where *V* denotes the set of nodes in the network (with *|V|* = *n*), *E* represents the edges on *V*, and *W* contains the weights on the edges. The goal of network enhancement is to generate a network *G*\* = (*E*\*, *V*, *W*\*) that provides a better representation of the underlying module membership than the original network *G*. For the analysis below, we let *W* represent a symmetric, non-negative matrix.

Diffusion-based models often rely on the row-normalized transition probability matrix *P* = *D*^−1^*W*, where *D* is a diagonal matrix whose entries are 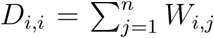. However, transition probability matrix *P* defined in this way is generally asymmetric and does not induce a directly usable node-node similarity metric. Additionally, most diffusion-based models lack spectral analysis of the denoised model. To construct our diffusion process and provide a theoretical analysis of our model, we propose to use a symmetric, doubly stochastic matrix (DSM). Given a matrix *M* ∈ ℝ^*n×n*^, *M* is a DSM if it satisfies the following conditions:

1. 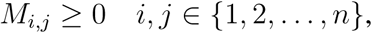
2. 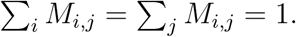

The second condition above is equivalent to **1** = (1, 1, …., 1)^*T*^ and **1**^*T*^ being a right and left eigenvector of *M* with eigenvalue 1. In fact, 1 is the greatest eigenvalue for all DSM matrices (see the remark following the definition of DSM in the Supplementary Notes) Overall, the DSM property imposes a strict constraint on the scale of the node similarities and provides a scale-free matrix that is well-suited for subsequent analyses.

### Network Enhancement (NE)

Given a matrix of edge weights *W* representing the pairwise weights between all the nodes, we construct another localized network *𝒯* ∈ ℝ^*n*×*n*^ on the same set of nodes, but we assign non-zero weights only to edges with high weights as follows. Denote the set of *K*-nearest neighbors (KNN) of the *i*-th node (including the node *i*) as *𝒩_i_*. We use these nearest neighbors to measure local affinity. Then the corresponding localized network *𝒯* can be constructed from the original weighted network using the following two steps:

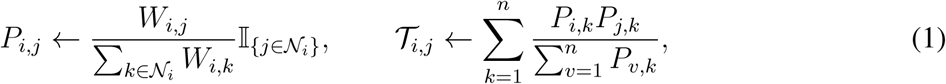

where 𝕀_{·}_ is the indicator function. We can verify that *𝒯* is a symmetric DSM by directly checking the conditions of the definition (**Supplementary Note 2**). *𝒯* encodes the local structures of the original network with the intuition that local neighbors (highly similar pairs of nodes) are more reliable than remote ones, and local structures can be propagated to non-local nodes through a diffusion process on the network. Motivated by the updates introduced in Zhou *et al.* ^32^, we define our diffusion process using *𝒯* as follows:

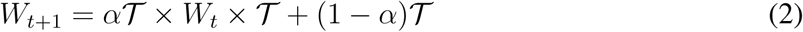

where *α* is a regularization parameter and *t* represents the iteration step. The value of *W*_0_ can be initialized to be the input matrix *W*. Eqn. (2) shows that diffusion process in NE is defined by random walks of length three or less and a form of regularized information flow. There are three main reasons for restricting the influence of random walks to at most third-order neighbors in the network: (1) for most nodes third-order neighborhood spans the extent of almost the entire biological network, making neighborhoods of order beyond three not very informative of individual nodes ^33, 34^, (2) currently there is little information about the extent of influence of a node (i.e., a biological entity, such as gene) on the activity (e.g., expression level) of its neighbor that is more than three hops away ^35^, and (3) recent studies have empirically demonstrated that network features extracted based on three-hop neighborhoods contain the most useful information for predictive modeling ^36^.

To further explore Eqn. (2) we can write the update rule for each entry:

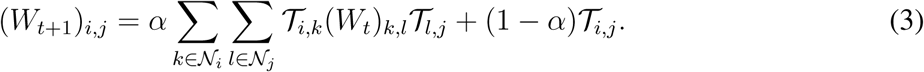

It can be seen from Eqn. (3) that the updated network comes from similarity/interaction flow going only through the neighbors of each data point. The parameter *α* adds strengths to self-similarities, i.e., a node is always most similar to itself. One key property NE that differentiates our method from typical diffusion methods is that in the local consistent diffusion process defined in Eqn. (2), for each iteration *t*, *W_t_* remains a symmetric DSM. Furthermore, *W_t_* converges to a non-trivial equilibrium network which is symmetric and a DSM as well **Supplementary Note 2**. This shows that Network Enhancement constructs an undirected network that preserves the DSM property of the original network. Through extensive experimentation, we show that NE improves the similarity between related nodes and the performance of downstream methods such as community detection algorithms.

The main theoretical insight into the operation of NE is that the proposed diffusion process does not change eigenvectors of the initial DSM while mapping eigenvalues via a non-linear function (**Supplementary Note 2**). Assume the initial symmetric DSM, *𝒯*_0_ has eigen-pair (*λ*_0_, **v**_0_), then the local consistent diffusion process defined in Eqn. (2) does not change the eigenvector and the final converged graph has eigen-pair (*f_α_*(*λ*_0_), **v**_0_), where 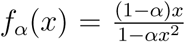. This property shows that, the diffusion process using a symmetric, doubly stochastic matrix is a non-linear operator on the spectrum of the eigenvalues of the original network. This results has a number of consequences. Practically, it provides us with a closed-form expression for the final converged network. Theoretically, it hints at how this diffusion process affects the eigen-spectrum and improves the network for subsequent analyses. 1) If the original eigenvalue is either 0 or 1, the diffusion process preserves this eigenvalue. This implies that, like other diffusion processes, NE does not connect disconnected components. 2) NE increases the gap between large eigenvalues of the original network and reduces the gap between small eigenvalues of this matrix. Larger eigengap is associated with better network community detection and higher-order network analysis ^12, 15, 16^. 3) The diffusion process always decreases the eigenvalues, which follows from: 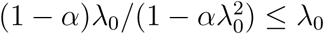, where smaller eigenvalues get reduced at a higher rate. This observation can be interpreted in relation to principal component analysis (PCA) where, in contrast to NE, the eigenspectrum below a threshold value is ignored. PCA has many attractive theoretical properties, especially for dimensionality reduction. In fact, Mashup ^17^, a feature learning method whose output is also a denoised version of the original network, can be fit by computing the PCA decomposition on the stationary state of the network. As Mashup aims to learn a low-dimensional representation of nodes in the network, the use of PCA is a natural choice for its objective. However, a smoothed-out version of the PCA is more attractive for network denoising because denoising is typically used as a preprocessing step for downstream prediction tasks, and thus robustness to the selection of a threshold value for the eigenspectrum is highly desirable.

These findings shed light on why the proposed algorithm (NE) enhances the robustness of the diffused network compared to the input network (**Supplementary Note 2**). In some contexts, we may need the output to remain a network of the same scale as the input network. This requirement can be satisfied by first recording the degree matrix of the input network and eventually map the denoised network back to the original scale by a symmetric matrix multiplication. We summarize our Network Enhancement algorithm along with this optional degree-mapping step in **Supplementary Note 2**.

### Code and Data Availability

All relevant data are public and available from the authors of original publications. The project website is at http://snap.stanford.edu/ne. The website contains preprocessed data used in the paper together with raw and enhanced networks. Source code of the NE method is available for download from the project website.

## Author information

The authors declare no conflict of interest. Correspondence should be addressed to J.L. and to S.B.

## Additional information

**Supplementary Information** contains a detailed description of datasets (**Supplementary Data**), additional experiments (**Supplementary Figures** and **Supplementary Note 1**), mathematical derivation, and theoretical analysis of NE (**Supplementary Note 2**).

